# IL-33 priming and antigenic stimulation synergistically promote the transcription of proinflammatory cytokine and chemokine genes in human skin mast cells

**DOI:** 10.1101/2022.04.14.488379

**Authors:** Junfeng Gao, Yapeng Li, Xiaoyu Guan, Zahraa Mohammed, Gregorio Gomez, Yvonne Hui, Dianzheng Zhao, Carole A. Oskeritzian, Hua Huang

## Abstract

**Background:** Antigenic stimulation through cross-linking the IgE receptor and epithelial cell-derived cytokine IL-33 are potent stimuli of mast cell (MC) activation. Moreover, IL-33 primes a variety of cell types, including MCs to respond more vigorously to external stimuli. However, target genes induced by the combined IL-33 priming and antigenic stimulation have not been investigated in human skin mast cells (HSMCs) in a genome-wide manner. Furthermore, epigenetic changes induced by the combined IL-33 priming and antigenic stimulation have not been evaluated.

**Results:** We found that IL-33 priming of HSMCs enhanced their capacity to promote transcriptional synergy of the *IL1B* and *CXCL8* genes by 16- and 3-fold, respectively, in response to combined IL-33 and antigen stimulation compared to without IL-33 priming. We identified the target genes in IL-33-primed HSMCs in response to the combined IL-33 and antigenic stimulation using RNA sequencing (RNA-seq). We found that the majority of genes synergistically upregulated in the IL-33-primed HSMCs in response to the combined IL-33 and antigenic stimulation were predominantly proinflammatory cytokine and chemokine genes. Moreover, the combined IL-33 priming and antigenic stimulation increase chromatin accessibility in the synergy target genes but not synergistically. Transcription factor binding motif analysis revealed more binding sites for NF-κB, AP-1, GABPA, and RAP1 in the induced or increased chromatin accessible regions of the synergy target genes.

**Conclusions:** Our study demonstrates that IL-33 priming greatly potentiates MCs’ ability to transcribe proinflammatory cytokine and chemokine genes in response to antigenic stimulation, shining light on how epithelial cell-derived cytokine IL-33 can cause exacerbation of skin MC-mediated allergic inflammation.

## Background

Mast cells (MCs) are long-lived tissue-resident cells with evolutionarily conserved functions in many inflammatory settings, including allergic inflammation and host defense against pathogen infection [1–3]. The release of these mediators initiates a cascade of immune responses, resulting in allergic reactions. MCs are dispersed throughout the boundaries between the external environment and tissues, such as around blood vessels, in the skin, and at mucosal surfaces. Because of their strategic locations, MCs are among the first exposed to antigenic stimulations and cytokines and chemokines secreted by injured epithelial cells and other inflammatory cells at the site of inflammation. MCs express FcεRI, the high-affinity receptor for IgE, which consists of one IgE-binding α subunit, one signaling β subunit, and a signaling γ subunit dimer. MCs can be activated by antigen (Ag) crosslinking of FcεRI-bound Ag-specific IgE (IgECL), which is produced in an adaptive immune [4]. Ag crosslinking of the IgE/FcεRI complex induces MC release of histamine, which is pre-made and stored in MC granules, and de novo gene transcription [5, 6].

Epithelial cell-derived cytokines IL-25, IL-33, and TSLP have been reported to activate MCs to amplify inflammation [7, 8]. IL-33 is a member of the IL-1 family [9, 10]. It is expressed in epithelial cells, dendritic cells, and macrophages in tissues such as the skin, colon, lymph nodes, and lung [11, 12]. Recent research has highlighted the role of interleukin-33 (IL-33) in modulating skin inflammatory cascades and its potential as a therapeutic target for atopic dermatitis [13, 14]. IL-33 protein is released from injured epithelial cells upon infections and inflammation as IL-33 stored in the nucleus of destroyed epithelial cells is set free [10, 15]. MCs express high levels of IL-33 (IL1RL1) receptor [16–18]. IL-33 activates MAP kinases and induces NF-κB phosphorylation [12]. IL-33 promotes the differentiation of MC progenitor cells into mature MCs that produce Th2 cytokines IL-5 and IL-13 [18]. Moreover, it directly activates MCs to produce pro-inflammatory cytokines and chemokines, including IL-6, IL-10, TNF, GM-CSF, CXCL8, and CCL1 [7, 19]. It synergizes with TSLP to promote MC differentiation and activation [20]. IL-33 also synergizes with IgE receptor activation of primary human MCs to promote cytokine and chemokine production without IL-33 priming [7, 21]. While previous studies have used qPCR to measure the synergistic effects of antigen and IL-33 stimulations on mRNA expression of *CCL1*, *CCL2*, *CXCL1*, *CXCL2*, *CXCL8*, *IL1B*, *IL5*, *IL6*, *IL13* genes in various MCs [7, 19, 21, 22], genome-wide profiling has not been conducted.

Moreover, the role of IL-33 priming in enhancing the transcription of cytokine and chemokine genes in MCs has not been clearly established. IL-33 primes a variety of cell types, including MCs to respond more vigorously to external stimuli [23, 24]. IL-33 has been shown to play a crucial role in promoting IgE-mediated allergic inflammation in vivo [25]. Notably, studies have demonstrated that IL-4, IL-5, and IL-13 levels are increased in ovalbumin (OVA)-sensitized mice treated with IL-33, compared to mice treated with IL-33 alone or OVA alone [26], suggesting that prolonged exposure to IL-33 can further increase MC responses to antigenic stimulation.

In this study, we sought to determine whether pretreatment with IL-33 for 24 hours can promote gene expression in response to the combined IL-33 and IgECL stimulation compared to without IL-33 pretreatment. We demonstrate that IL-33 priming of HSMCs enhanced their capacity to promote transcriptional synergy of the *IL1B* and *CXCL8* genes in response to antigen stimulation compared to without IL-33 priming. We identified the target genes in the IL-33-primed HSMCs upon combined IL-33 and antigenic stimulation using RNA sequencing (RNA-seq). We found that the majority of genes synergistically upregulated in the IL-33-primed HSMCs in response to the combined IL-33 and antigen stimulation were predominantly proinflammatory cytokine and chemokine genes. Moreover, the combined IL-33 and antigen stimulation increased chromatin accessibility in the synergy genes but not synergistically in the IL-33-primed HSMCs. Our study shines a light on a mechanism by which IL-33 might cause exacerbation of skin MC-mediated allergic inflammation.

## Results

### The combined IL-33 priming and antigenic stimulation promote expression of two clusters of genes

To investigate the synergistic gene transcription promoted by antigen and epithelial cell-derived cytokines IL-33 in human skin-derived mast cells (HSMCs), we first compared whether pretreatment with IL-33 for 24 hours can promote gene expression in response to the combined IL-33 and IgECL stimulation compared to without IL-33 pretreatment. We chose to measure *IL1B* and *CXCL8* mRNA expression, which is known to be upregulated by the combined IgECL and IL-33 stimulation [7, 19–22] by qPCR. We used a published method to analyze transcriptional synergy, which is defined transcriptional synergy as greater than additive effects and is calculated by the formula synergistic log_2_ [*E*_IgECL+IL-33_/(*E*_IgECL_+*E*_IL-33_)] [7, 27, 28]. Specifically, log2FC>0 indicates positive transcriptional synergy, while log_2_FC<0 represents negative transcriptional synergy. A log_2_FC=0 indicates additive. We found that IL-33 pretreatment for 24 hours promoted transcriptional synergy induced by the combined IL-33 and IgECL stimulation (log_2_FC *IL1B*=3.5, *CXCL8*=2.9), resulting in 16- and 3-fold increase in transcriptional synergy for *IL1B* and *CXCL8,* respectively, compared to without IL-33 pretreatment (log_2_FC *IL1B*=1.5, *CXCL8*=1.9) (Fig. 1). Statistical differences among the single and combined treatments were analyzed using two-way ANOVA as recommended by the published paper [29]. Our data demonstrate that although with or without IL-33 pretreatment, IgECL and IL-33 stimulation can both promote transcriptional synergy for the *IL1B* and *CXCL8* genes, pretreatment with IL-33 dramatically enhances the transcriptional synergy.

**Figure 1.**
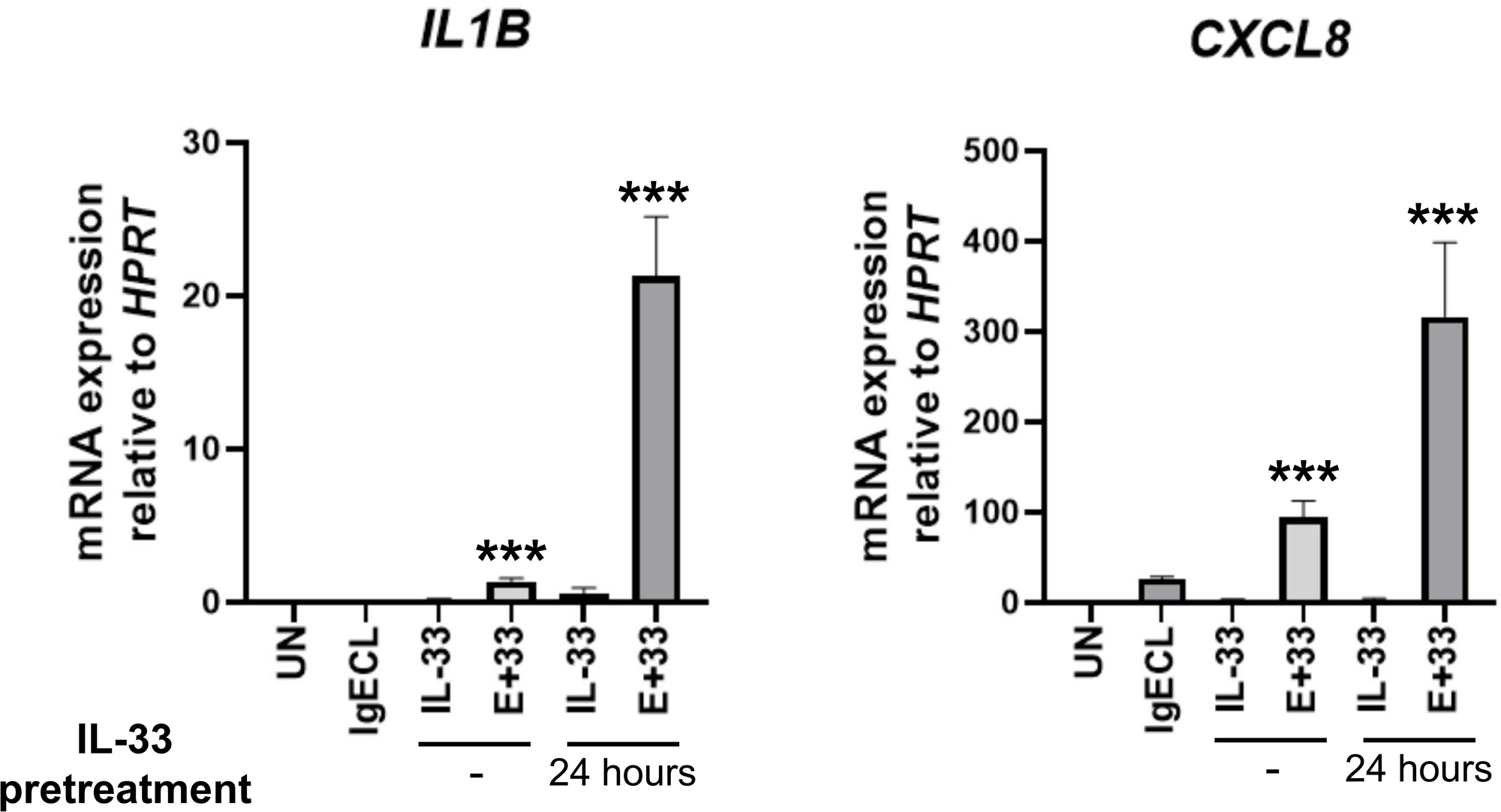
IL-33 priming and antigenic stimulation synergistically promote the transcription of *IL1B* and *CXCL8* genes in HSMCs. HSMCs were not treated (UN) or treated with IgE receptor cross-linking (IgECL) for 2 hours, IL-33 for 2 hours or 26 hours (pre-treated for 24 hours prior to the addition of the other stimuli treatment for 2 hours), or in combination of IgECL and IL-33 (2 hours or 26 hours). qPCR analysis of the mRNA expression of *IL1B* and *CXCL8*. Statistical differences were analyzed using two-way ANOVA. Data represents mean ± SEM of four biological samples. *** *P* < 0.001.

Given that many molecular events can occur during the 24-hour pretreatment, we thus referred to the pretreatment with IL-25, TSLP, or IL-33 for 24 hours in the combined stimulations as priming. For the combined treatments, we pretreated HSMCs with cytokines IL-25, TSLP, also produced by epithelial cells, or IL-33 for 24 hours before subjecting them to the combined cytokines and IgECL stimulation for an additional 2 hours. We found that IgECL stimulation alone upregulated 501 genes with increased expression greater than 2-fold and 37 genes with increased expression greater than 10-fold (Fig. 2A, 3A and Table S1, sheet 1). We found that IL-33 priming alone upregulated 259 genes that were expressed 2-fold greater and 22 genes with increased expression greater than 10-fold (Fig. 2B, 3A and Table S1, sheet 2). Whereas IL-25 priming alone only upregulated nine genes greater than 2-fold with none of the genes increased expression greater than 10-fold (Fig. 2C, 3A and Table S1, sheet 3). TSLP upregulated 17 genes greater than 2-fold with one gene increased expression greater than 10-fold (Fig. 2D, 3A and Table S1, sheet 4). These results indicate that IL-33 is a more potent stimulus than IL-25 or TSLP in promoting gene expression in HSMCs.

**Figure 2.**
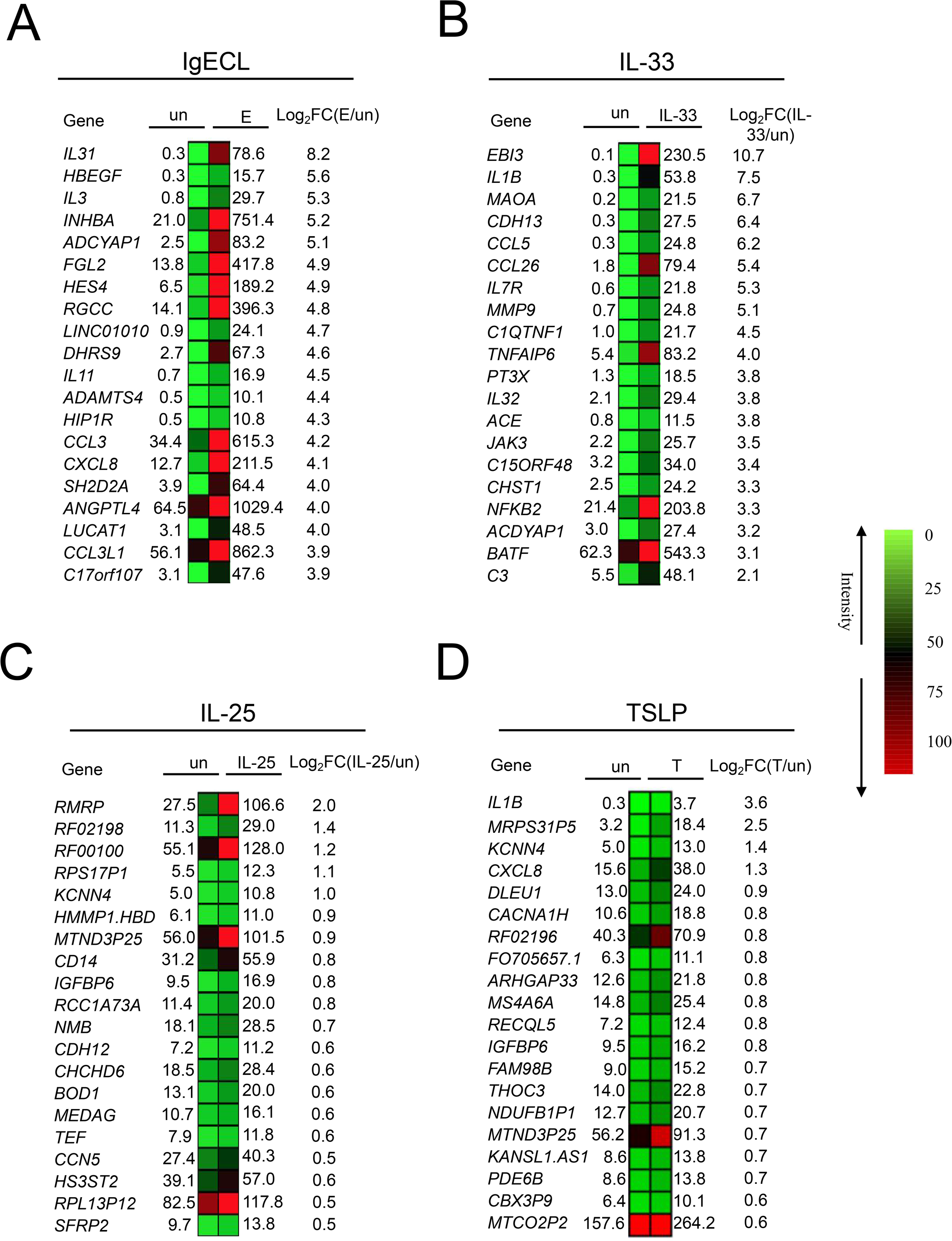
Differential mRNA expression analysis of RNA-seq prepared from HSMCs that were exposed to various treatments. Heatmap representations of top ranked genes after IgE receptor cross-linking treatment alone (A), IL-33 treatment alone (B), IL-25 treatment alone (C), or TSLP treatment alone (D). Legends: un, not treated; E, IgECL; IL-33, IL-33 treatment; IL-25, IL-25 treatment; T, TSLP treatment; log_2_FC, log_2_ fold change. The numbers indicate RNA reads (RPKM). Data A-D represent two biological samples.

**Figure 3.**
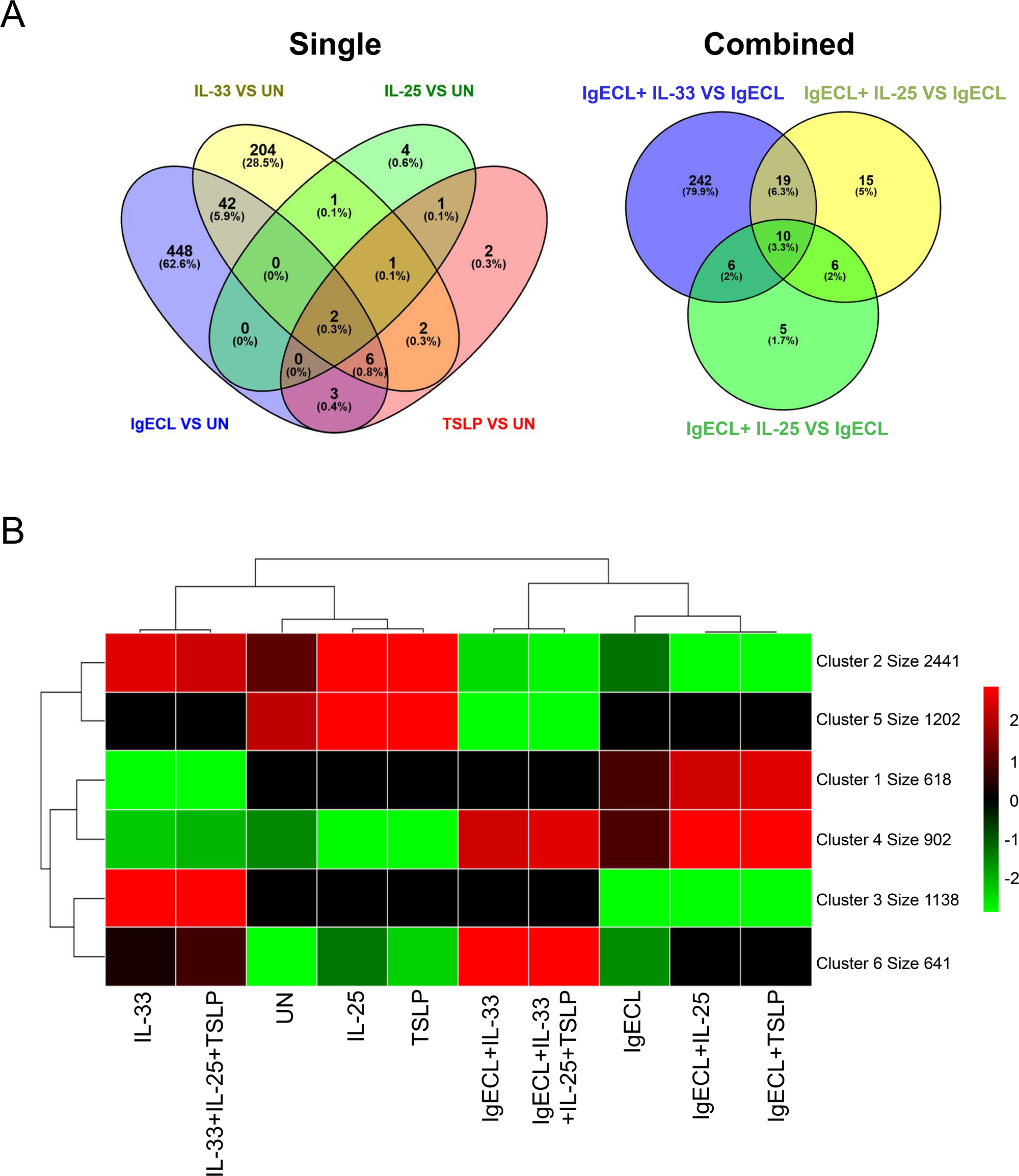
IL-33 priming acts synergically with IgECL to promote gene transcription in clusters four and six. HSMCs were not treated (UN), treated with IgE receptor cross-linking (IgECL) for 2 hours, IL-33 for 26 hours (priming for 24 hours prior to the addition of the other stimuli treatment for 2 hours), IL-25 for 26 hours, TSLP for 26 hours alone (single) or in combinations (combined) as indicated. (**A**) Venn diagram showing the overlap of upregulated differentially expressed genes among the four comparisons of IL-33 vs UN, IL-25 vs UN, IgECL vs UN, TSLP vs UN (left panel), and the comparisons of IgECL+ IL-33 vs IgECL, IgECL+ IL-25 vs IgECL, IgECL+ TSLP vs IgECL (right panel). (**B**) K-means clustering analysis was performed using R package pheatmap. The cluster size numbers indicate the number of genes that are grouped in the cluster. RNA-seq data were generated from two biological samples.

To determine whether any of the combinatorial stimulations may also upregulate HSMC genes, we used the K mean clustering method to group genes into clusters with similar expressions. We first determined the optimal K numbers of six using Elbow Plot (Fig. S1). Heatmap K means clustering analysis revealed that the combined IL-33 and antigen treatment in the IL-33-primed HSMCs upregulated the expression of two major clusters of genes (cluster four and cluster six) relative to IgECL or IL-33 single stimulation (Fig. 3B). There were 902 genes in cluster four and 641 genes in cluster six (Table S2). IL-25 and TSLP, when added with IgECL, also promoted expression of genes in cluster four. However, the magnitude of upregulation induced by IL-25 and TSLP was lower compared to IL-33. IL-25 and IgECL upregulated 50 genes compared with IgECL alone (Fig. 3A, S2A and Table S3, sheet 1). TSLP and IgECL upregulated 27 genes compared with IgECL alone (Fig. 3A, S2B and Table S3, sheet 2). The addition of IL-25 and TSLP to the IgECL and IL-33 combination did not significantly alter MC gene expression profiles compared with the IgECL and IL-33 combination (Fig. 3A, S2C and Table S3, sheet3). These results demonstrated that epithelial cell-derived cytokines IL-33 together with IgECL promoted the expressions of cluster four and six HSMC genes.

### The combined IL-33 and antigen stimulation in the IL-33-primed HSMCs synergistically induce pro-inflammatory gene transcription

We performed gene ontology (GO) analysis on genes in clusters four and six to determine which types of genes were enriched among genes that were promoted by the combined IL-33 and antigen stimulation in the IL-33-primed HSMCs. We found that genes encoding for cytokines (Cluster four, p=1.0×10^−5^; cluster six, p=2.5×10^−5^, Fig. 4A), receptors and receptor binding (Cluster four, p=7.9×10^−2^; cluster six, p=3.6×10^−2^, Fig. 4A), signaling molecules (Cluster four, p=9.7×10^−6^; cluster six, p=7.3×10^−3^, Fig. 4A), transcription factors (Cluster four, p=1.8×10^−2^; cluster six, p=2.6×10^−3^, Fig. 4A), NF-κB signaling pathway (Cluster four: p=6.5×10^−2^, cluster six: p=4.2×10^−2^, Fig. 4A), chemokines (Cluster four: p=1.1×10^−5^; cluster six, p=9.7×10^−2^, Fig. 4A), and molecules involved in cell death (Cluster four, p=2.6×10^−2^; cluster six: p=1.7×10^−2^, Fig. 4A) were significantly enriched in both clusters.

**Figure 4.**
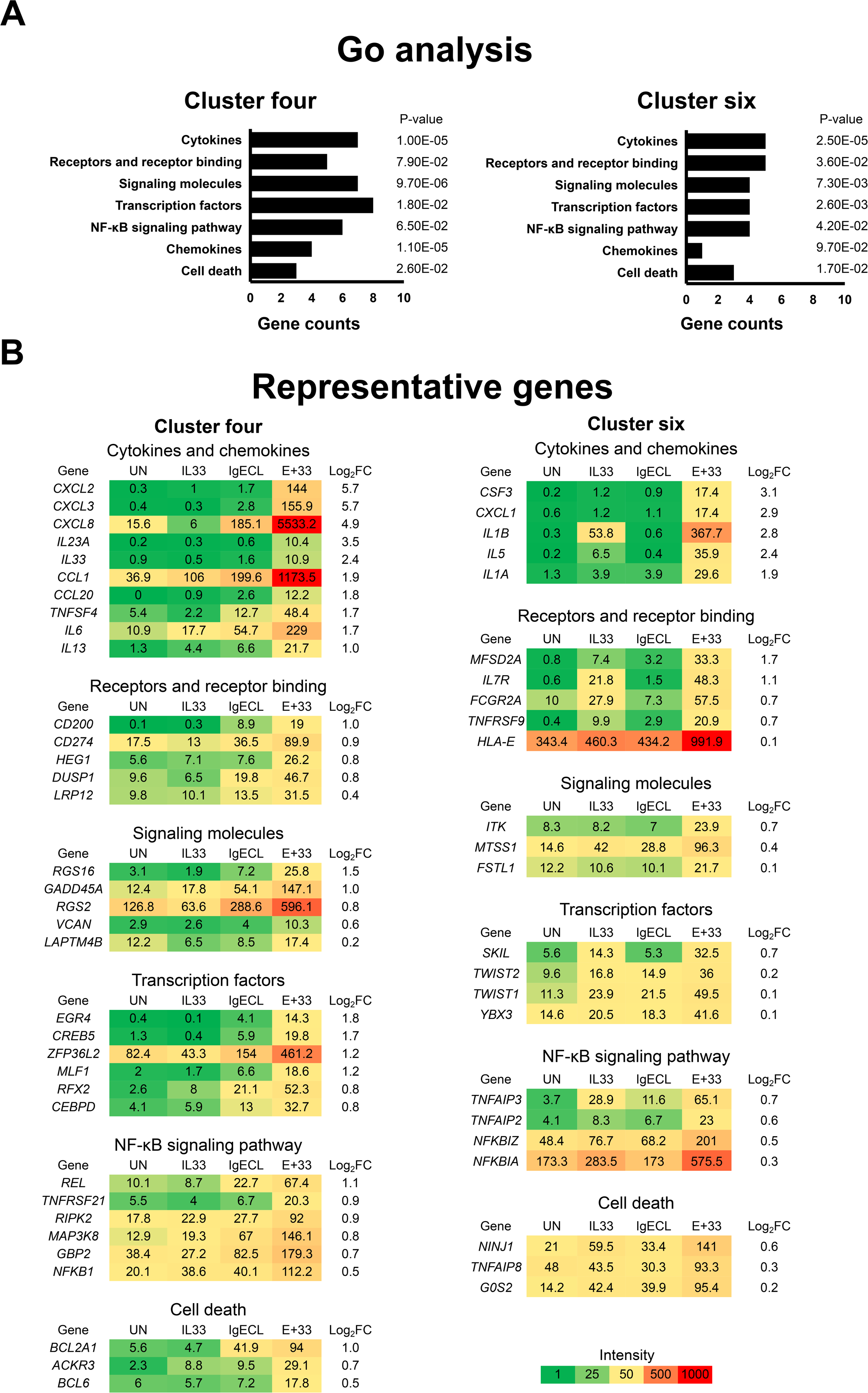
Genes synergically upregulated by IL-33 priming and IgE receptor cross-linking are enriched in genes encoding pro-inflammatory cytokines and chemokines. (**A**) Gene ontology (GO) enrichment analysis of the upregulated genes in clusters four and six. *P*-values were calculated by a one-side Fisher’s exact test with the adjustment of Benjamini-Hochberg method. (**B**) Heatmap representation of top ranked genes in each gene family in clusters four and six. The numbers indicate RNA reads (FPKM). log_2_FC: log_2_[*E*_IgECL+IL-33_/(*E*_IgECL_+*E*_IL-33_)]. Data represents the average reads generated from two biological samples.

The top-ranked representative genes in clusters four and six are listed in Fig. 4B. Genes encoding cytokines and chemokines family in cluster four were most significantly enriched. They include *CXCL8*, *CXCL3*, *TNFSF4*, *IL33*, *IL6*, *CCL20*, *IL23A*, *CCL1*, *CXCL2*, and *IL13* (Fig. 4B). Genes enriched in the cytokines and chemokines gene family in cluster six include *IL1B*, *IL5*, *CSF3*, *CXCL1* and *IL1A.* While *CXCL8*, *IL5*, *IL13*, and *IL6* have been reported to be upregulated synergistically by the combined IL-33 and antigen stimulation [22, 30, 31], *TNFSF4*, a member of TNF family, *CXCL3*, *IL33*, *CCL20*, *IL23A*, *CCL1*, *CXCL2*, *CXCL1*, *CSF3*, and *IL1A*, have not been previously reported as target genes of the combined IL-33 and antigen stimulation in HSMCs.

Genes encoding signaling molecules in cluster four were also highly enriched (Fig. 4A). These genes include *RGS2*, which potently interferes with signaling through receptors that couple to Gαq [32], and *RGS16*, which inhibits signal transduction through the Gα13-RHO [33]. *MTSS1* and *ITK* were enriched in cluster six (Fig. 4B). These genes were not reported previously as target genes for the synergistic stimulation of IL-33 and antigen. We also found that genes encoding receptor and receptor binding molecules were significantly enriched in clusters four and six (Fig. 4A). These genes included *IL7R*, a receptor for IL-7 and TSLP [34–36], *FCGR2A,* which encodes a cell surface receptor belonging to the family of immunoglobulin Fc receptor, and *CD274*, encoding PD-L1, a ligand that binds to the receptor PD1 commonly found on T-cells to block T-cell activation [37, 38] (Fig. 4B).

Many genes encoding molecules involved in cell survival and cell death were identified in clusters four and six (Fig. 4A). *BCL2A1* is a highly regulated NF-κB target gene that exerts important pro-survival functions [39]. BCL6 acts to prevent apoptotic cell death in differentiating myocytes [40]. *ACKR3*, which encodes a receptor for CXCL12 and CXCL11 [41], has been reported to inhibit apoptosis [42]. TNF Alpha Induced Protein 8 (TNFAIP8) was showed to increase cell survival and decrease apoptosis in hepatocellular carcinoma cells [43]. On the other hand, we also found that IL-33 and IgECL stimulation upregulated genes involved in apoptosis, including *G0S2,* which promotes apoptosis by binding to BCL2. *NINJ1,* which encodes NINJURIN-1 [44], has been shown to mediate plasma membrane rupture during lytic cell death [45] and promote apoptosis in macrophages during early ocular development [46].

Genes encoding transcription factors were found to be significantly enriched in clusters four and six (Fig. 4A). EGR4, activated by NFAT in response to antigenic stimulation, regulates pro-inflammatory cytokine gene expression [47], whereas CEBPD results in increased *IL6* gene transcription [48]. Transcription factors *CREB5*, *MLF1*, *ZFP36L2* and *RFX2* were enriched in cluster four, while transcription factors *SKIL*, *TWIST2*, *YBX3* and *TWIST1* were enriched in cluster six (Fig. 4B). Genes encoding NF-κB signaling pathway were also highly enriched in both clusters four and six (Fig. 4A). These genes included *NFKB1*, encoding p50 subunit of NF-κB transcription factor complex that can lead to amplified transcription outputs for many of the NF-κB target genes, such as *IL1B*, *IL6*, *CXCL8*, *TNF*, *MCP1*, and *CXCL1* [49]. *REL*, *TNFRSF21*, *MAP3K8*, *GBP2* and *RIPK2* were enriched in cluster four, while *TNFAIP3*, *NFKBIA*, *NFKBIZ* and *TNFAIP2* were enriched in cluster six (Fig. 4B). Our results demonstrate that proinflammatory cytokine and chemokine genes were significantly enriched among genes highly responsive to the combined IL-33 and antigen stimulation in the IL-33-primed HSMCs as those genes with log_2_FC>1.0.

RNA-seq results were validated by performing a qPCR analysis on selected genes whose products can promote MC survival, form positive feed-forward loops, or enable MCs to perform new functions using HSMCs derived from four independent donors. We confirmed the transcriptional synergy for *CXCL2* (log_2_FC=3.9), *CXCL8* (log_2_FC=1.9), *IL1B* (log_2_FC=2.0), *CCL1* (log_2_FC=1.4), *NAMPT* (log_2_FC=1.6), *IL7R* (log_2_FC=0.9), *CD274* (log_2_FC=0.3), *FCGR2A* (log_2_FC=0.3), *NFKB1* (log_2_FC=0.1) and *TNFAIP8* (log_2_FC=0.5) in the IL-33-primed HSMCs in response to the combined IL-33 and IgECL stimulation (Fig. 5A). Consistent with the transcriptional synergy, synergy at the protein level was also detected for IL1B (log_2_FC=1.9) and CXCL8 (log_2_FC=3.1) (Fig. 5B). CXCL2 protein expression was also measured but was not detectable using the commercial CXCL2 ELISA kit.

**Figure 5.**
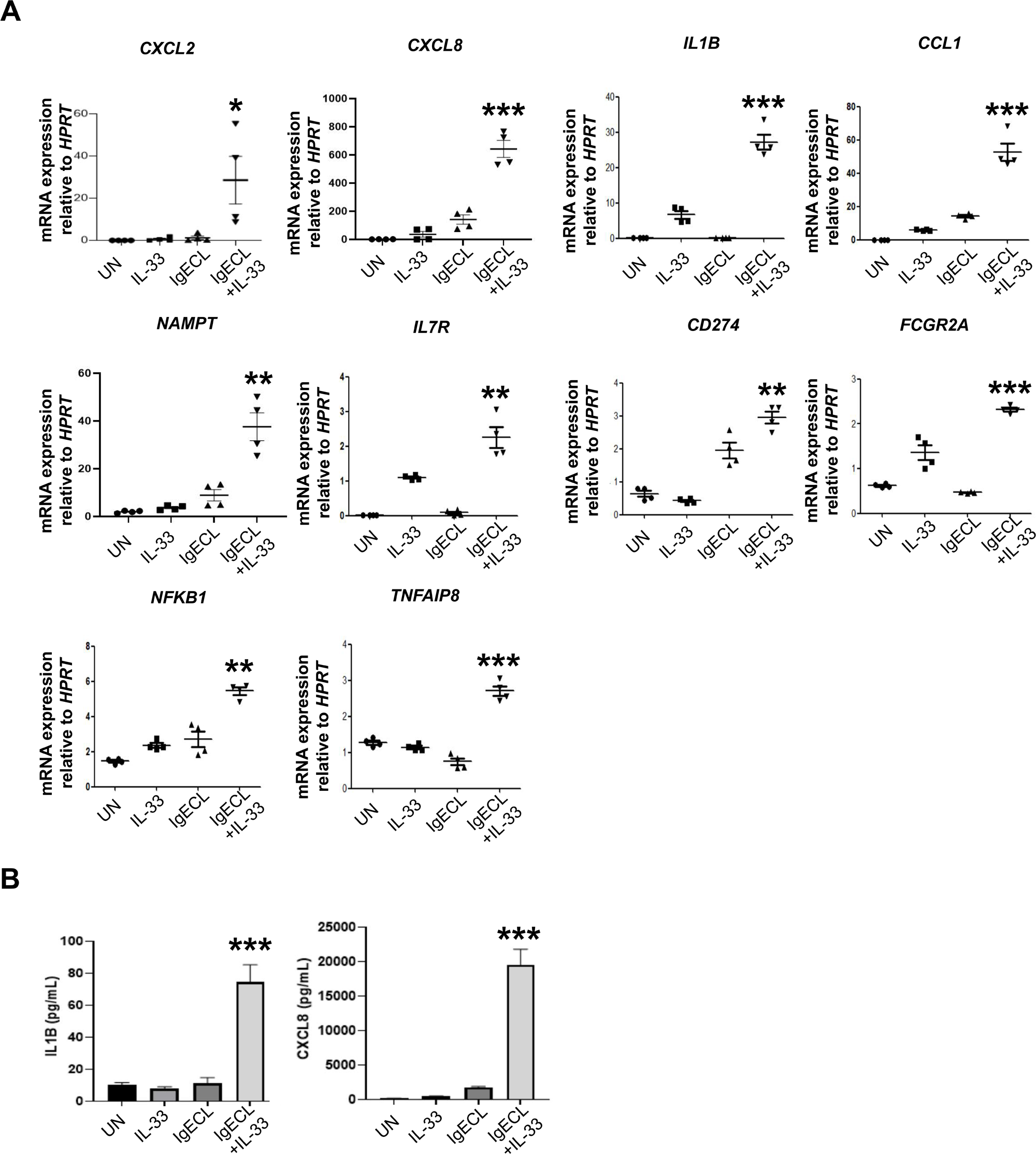
Validation of gene expression promoted by the combined IL-33 priming and IgE receptor cross-linking. HSMCs were not stimulated (UN) or stimulated with IgE receptor cross-linking (IgECL) for 2 hours, IL-33 for 26 hours (priming for 24 hours prior to the addition of the other stimuli treatment for 2 hours), or in combination of IgECL and IL-33. (**A**) qPCR analysis of the mRNA expression of activation-induced genes identified by RNA-seq. (**B**) The same treatments as described under (A) were applied, supernatants were collected after 24 hours and IL-1B and CXCL8 were quantified by ELISA. Statistical differences were analyzed using two-way ANOVA. Data represents mean ± SEM of four biological samples. * *P* < 0.05, ** *P* < 0.01, *** *P* < 0.001.

### The combined IL-33 and antigen stimulation increases chromatin accessibility in the synergy genes but not synergistically in the IL-33-primed HSMCs

Enhancers integrate signal inputs and convert them into transcriptional outputs [50]. Thus, it is likely that the combined IL-33 and antigen treatment activate signal-dependent transcription factors that, in turn, bind to enhancers to increase their chromatin accessibility in the IL-33-primed HSMCs. To investigate this, we performed Omni-ATAC-seq analysis on the HSMCs that were primed with IL-33 for 24 hours and treated with the combined IL-33 and IgECL stimulation for 2 hours or HSMCs that were treated IL-33 alone for 26 hours or IgECL alone for 2 hours. We used differential bind analysis (Diffbind) to compare ATAC-seq narrow peaks from the IL-33-primed HSMCs treated with the combined IL-33 and IgECL stimulation or from HSMCs that were treated with either IL-33 or IgECL alone. We assigned ATAC-seq peaks located within 100 kb upstream or downstream of the transcription start site (TSS) of a gene using Binding and Expression Target Analysis (BETA) [51]. Tens of thousands of narrow peaks (21,451), which were significantly enriched peaks compared to pooled and normalized background, increased significantly in the IL-33-primed HSMCs treated with the combined IL-33 and IgECL stimulation compared to IL-33 stimulated alone (Table S4). Fewer narrow peaks (521) were found to be significantly elevated when compared with IgECL stimulated alone (Table S4).

We analyzed the differences in the number of upregulated narrow peaks in positive transcriptional synergy genes (referred to as synergy target genes, with log2FC>0) and non-positive transcriptional synergy genes (referred to as non-synergy target genes, with log2FC≤0, Table S5). A significant genome-wide difference in upregulated narrow peaks was observed between synergy target genes and non-synergy target gene in response to IL-33 priming and antigenic stimulation (Fig. 6A). In Fig. 6B-D, we focused our analysis on the top-ranked *CXCL2*, *CXCL8*, and *IL1B* genes and assessed the normalized values of differentially increased ATAC-seq narrow peaks in the samples primed with IL-33 and treated with combined IL-33 and IgECL stimulation compared to those treated with either IgECL or IL-33 alone. We observed that the combined IL-33 and IgECL treatment led to increased chromatin accessibility, as indicated by higher narrow peak values at several candidate enhancers of the synergy target genes. Significantly increased narrow peak values were found for *CXCL2* gene at E+65 [IgE+IL-33/IL-33 (E+33/33) ratio=1.6], E+42 (E+33/33=1.5), PE (E+33/33=1.1), and E-10 (E+33/33=1.5) compared to IL-33 priming alone (Fig. 6B). Additionally, combined IL-33 and IgECL treatment led to significantly increased narrow peak values at E-35 (E+33/33=1.3), E-18 (E+33/33=1.5), E-5 (E+33/33=2.0), PE (E+33/33=1.3), and E+3 (E+33/33=1.4) of the *CXCL8* gene, compared to IL-33 priming alone (Fig. 6C). Furthermore, significantly increased narrow peak values were observed at E-18 (E+33/E=1.3), PE (E+33/E=1.3), and E+3 (E+33/E=1.5) of the *CXCL8* gene when compared to IgECL stimulation alone (Fig. 6C). In the *IL1B* gene, significant increases in narrow peak values were found at E+86 (E+33/33=1.6), E+71 (E+33/33=1.4), E+55 (E+33/33=1.5), E-3 (E+33/33=1.5), and E-43 (E+33/33=1.4) when compared to the IL-33 priming alone (Fig. 6D). However, no differential increased narrow peaks were observed for *CXCL2* and *IL1B* genes in comparison to IgECL alone (Fig. 6B, D).

**Figure 6.**
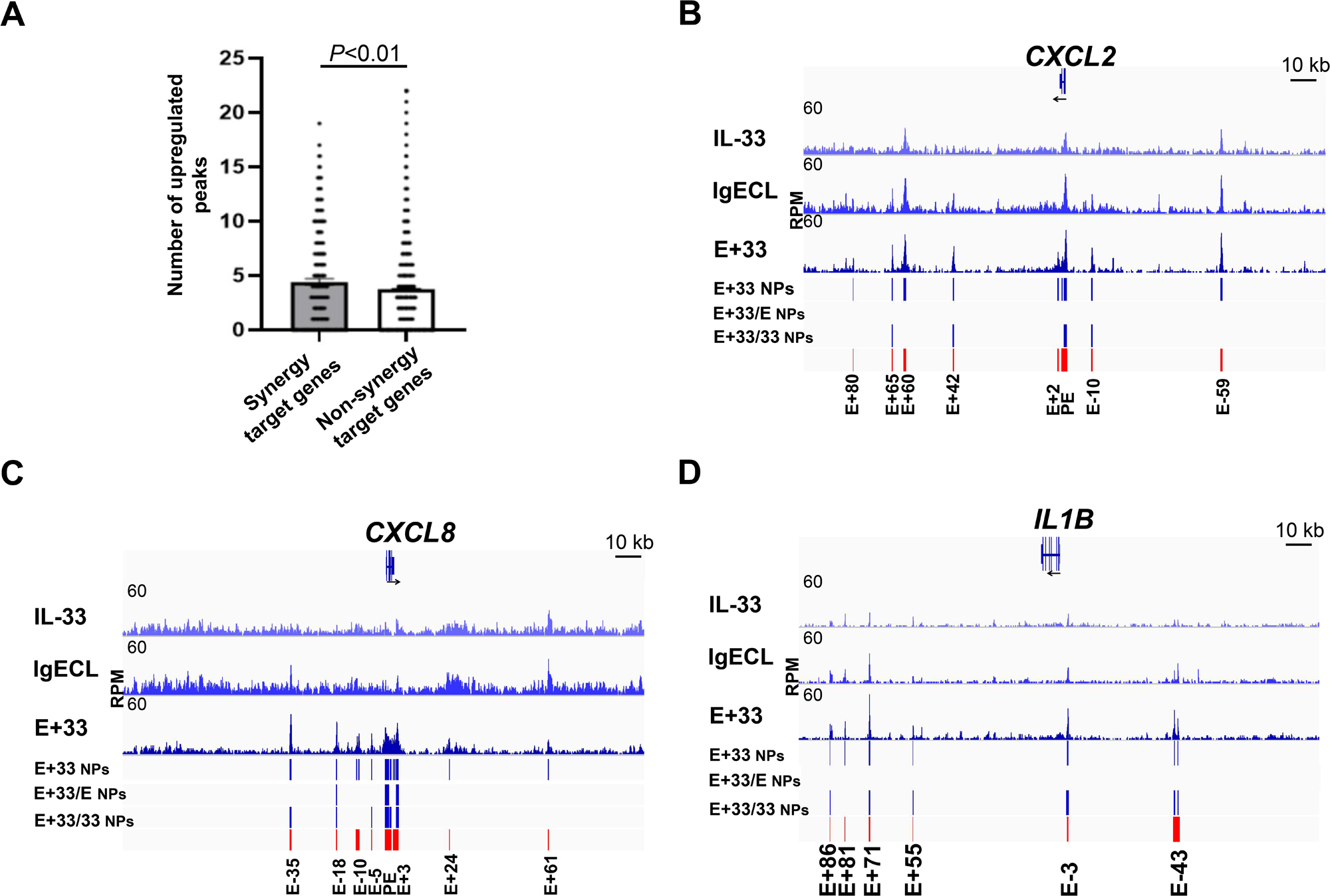
IL-33 priming and IgECL increase chromatin accessibility in the synergy target genes. (**A**) The numbers of upregulated ATAC-seq narrow peaks in the combined IL-33 priming and antigenic stimulation relative to IgECL or IL-33 stimulation were subtracted using Diffbind analysis. ATAC-seq was performed on HSMCs under various conditions, IgECL, IL-33 or IgECL plus IL-33. ATAC-seq narrow peaks (NPs) of *CXCL2* (**B**), *CXCL8* (**C**), *IL1B* (**D**) genes in HSMCs treated with IgECL and IL-33 (E+33) are shown (blue bars). Differential NPs generated using DiffBind comparing IgECL+IL-33 with IgECL (E+33/E NPs) or comparing IgECL+IL-33 with IL-33 (E+33/33 NPs) are also shown (blue bars). Candidate enhancers are marked with red bars and E - or + numbers, indicating distances (kb) from the TSS of the indicated genes. RPM, reads per million nucleotides mapped. The IGV tracks are generated from one biological sample, representing two biological replicates with similar patterns.

Only E-18, PE, and E+3 enhancers of the *CXCL8* showed increased narrow peaks in HSMCs treated with combined IL-33 and IgECL when compared to either IL-33 treatment or IgECL stimulation alone. The E+33/33 ratios were 1.5, 1.3, 1.4, for E-18, PE, E+3, respectively. Additionally, the E+33/E ratios were 1.3, 1.3, 1.5 for E-18, PE, E+3, respectively (Fig. 6C). These results indicate that the combined IL-33 priming and IgECL stimulation resulted in increased ATAC-seq peak values at certain enhancers of the synergy genes *CXCL2*, *CXCL8*, and *IL1B* in HSMCs but were less than additive compared to IL-33 priming or IgECL stimulation alone.

We analyzed transcription factor binding motifs in the chromatin accessible areas that were upregulated by the combined IL-33 and antigen stimulation in the IL-33-primed HSMCs relative to IgECL stimulation or IL-33 treatment alone using MEME-ChIP [52]. We found that NF-κB, AP-1, RAP1, and GABPA binding motifs were significantly enriched in the increased narrow peaks (Fig. 7). Our results suggest that genes responsiveness to the combined IL-33 and antigen stimulation in the IL-33-primed HSMCs is predetermined by the transcriptional codes embedded in the gene sequences. Furthermore, the combined IgECL and IL-33 treatment might achieve greater synergy in promoting gene transcription of the synergy target genes by binding to NF-κB, AP-1, GABPA and RAP1 binding sites.

**Figure 7.**
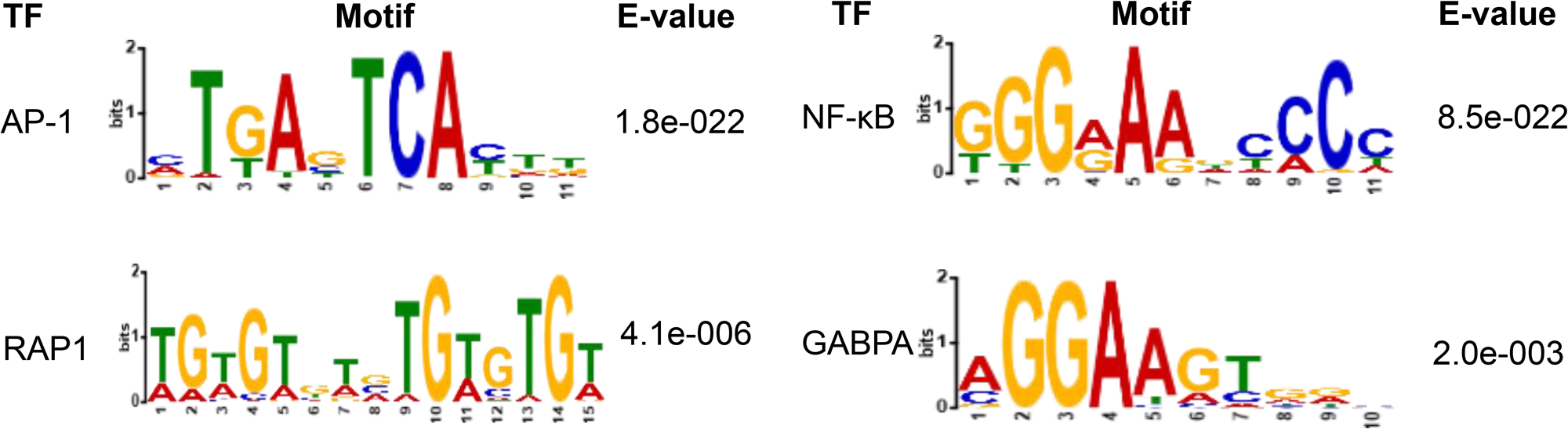
The AP-1, NF-κB, RAP1, and GABPA binding motifs are enriched in the regions with increased chromatin accessibility in the IL-33 priming and IgE receptor cross-linking-upregulated genes. Data represents two biological samples.

## Discussion

The results of this study provide direct evidence that IL-33 primes HSMCs with synergically enhanced capacity to respond to the combined IL-33 and antigen treatment. IL-33 is known to prime variety of cell types to respond more vigorously to external stimuli [23, 24]. It has been reported that IL-33 primes mouse MCs with amplified responses to ATP and IgG-mediated activation [53, 54]. Our study adds a new IL-33 priming capacity in HSMCs to the existing list. Compared with previous work that used a 15-minute IL-33 pretreatment [7], our work is designed to directly assess the effect of a 24 hour of IL-33 pretreatment on the upregulation of genes in HSMCs using a genome-wide and unbiased method. We demonstrate that IL-33 priming resulted in a 16-fold increase in transcriptional synergy for *IL1B* and a 3-fold increase for *CXCL8*. Similar to findings by the previous report [7], we confirmed a remarkable synergy at the protein level for CXCL8. Mechanisms by which this transcriptional synergy is not yet clear. A previous study used rat basophilic leukemia MCs, RBL-2H3 to demonstrate that multiple copies of NF-κB and NFAT binding sites were critical in antigen and IL-33 stimulation-driven reporter gene transcription. Our transcription factor motif analysis on chromatin accessible regions that were promoted by IL-33 priming and the subsequent combined IL-33 and antigen stimulation indeed established significantly enriched NF-κB binding motif and AP-1 binding motif, supporting previous conclusion that IL-33 and antigen stimulations synergistically activate NF-κB and NFAT to promote gene transcription of pro-inflammatory cytokines and chemokines *TNF*, *IL6*, *IL13*, *CCL2* (*MCP1*), *CCL7* (*MCP3*) and *CCL3* (*MIP1A*) [22]. However, NF-κB and NFAT are signal-dependent transcription factors. Activation of these transcription factors is rapid within minutes of receptor binding [55]. Therefore, the previous finding does not offer a complete explanation of IL-33 priming action. One best-studied example of cytokine priming is IL-4 priming for Th2 cell differentiation. IL-4 priming induces upregulation of GATA3, which then reprograms Th2 cells to acquire the capacity to transcribe the *Il4* gene [56]. It is possible a major transcription factor that depend on NF-κB and NFAT can be upregulated by IL-33 priming in HSMCs, conferring dramatically enhanced capacity to transcribe the cytokine and chemokine genes in the later response to antigenic stimulation or the combined IL-33 and antigen stimulation.

Our transcription factor motif analysis on IL-33 priming and antigenic stimulation responsive enhancers also provide some clues to the candidate transcription factors that are induced by IL-33 priming. Given that *NFKB1* (also known as p50) and *REL* (also known as p68) are the two subunits of NF-κB transcription factor complex and that GBP2, a guanine-binding protein, RIPK2, a member of the receptor-interacting protein (RIP) family of serine/threonine protein kinases, are known potent activator of the NF-κB pathway, the putative transcription factor whose induction dependent on NF-κB signaling can form a positive feed-forward loops to enhance HSMCs’ capacity to respond a combined secondary antigenic or the combined IL-33 and antigen stimulation. Interestingly, we also found potential negative feedback loops formed by TNFAIP2, TNFAIP3, NFKB1A, and NFKB1Z whose coding genes were significantly increased by the combined IL-33 and antigen stimulation. Therefore, upregulation of these gene transcriptions might provide critical control of NF-κB activation.

In addition to NF-κB binding motif and AP-1 binding motifs, we found more binding sites in the chromatin accessible regions that were promoted by combined IL-33 priming and antigen stimulation for GABPA, which binds to GA repeats and is a paralog gene of ETS2. While GABPA mRNA was expressed at low level revealed by our RNA-seq analysis, several ETS2 family transcription factor genes *ETS2*, *ELK1*, *ELF1*, *SPI1,* whose gene products can bind to GABPA binding site, were expressed in HSMCs. Whether these ETS2 family transcription factors can be recruited to bind GABPA binding sites in response to the combined IL-33 priming and antigenic stimulation remains to be determined.

Enhancers integrate signal inputs through recruiting signal-dependent transcription factor to their cognate transcription factor binding sites in the enhancers. Binding of transcription factors then recruit histone modifying enzymes to change H3K27 acetylation, leading to changes in chromatin accessibility [50, 57, 58]. We found IL-33 priming and the subsequent combined IL-33 and antigen stimulation indeed increased chromatin accessibility at the enhancers of synergy target genes. However, the chromatin accessibility increases induced by these processes did not reach additive or synergistic levels compared to IL-33 priming or antigen stimulation alone. These results suggest that the degree of changes in chromatin accessibility at individual enhancers might not correlate with the degree of transcriptional synergy. Further research deciphering transcriptional codes contained in the IL-33 priming responsive enhancers of synergy target genes could provide insights into the mechanisms underlying IL-33 priming for antigen-mediated MC activation and potentially lead to the development of targeted therapies for infection-exacerbation of allergic inflammation.

### Conclusions

In conclusion, our study provides evidence that IL-33 and IgECL stimulation can promote transcriptional synergy for mast cell genes, and IL-33 priming significantly enhances this synergy. We also found that combined IL-33 and antigen stimulation lead to the upregulation of two gene clusters in IL-33-primed HSMCs, cluster four (902 genes) and cluster six (641 genes), as indicated by heatmap K means clustering analysis. We demonstrate that proinflammatory cytokine and chemokine genes were significantly enriched among genes highly responsive to the combined IL-33 and antigen stimulation in the IL-33-primed HSMCs. Furthermore, analysis of Chromatin Accessibility revealed that although the combined IL-33 and antigen treatment in the IL-33-primed HSMCs amplifies chromatin accessibility in top-ranked *CXCL2*, *CXCL8* and *IL1B* genes, albeit the enhancement is not synergistic in nature. Overall, our study sheds light on the mechanism by which the epithelial cell-derived cytokine IL-33 may exacerbate skin MC-mediated allergic inflammation.

## Materials and Methods

### Human skin mast cell (HSMC) culture

De-identified fresh skin biopsies were obtained after breast reduction or mastectomy from the Cooperative Human Tissue Network of the National Cancer Institute. Handling human skin tissues was approved by the University of South Carolina Institutional Review Board. Human skin MCs were cultured from human skin essentially as described [59, 60]. After removing subcutaneous fat by blunt dissection, residual tissue was cut into 1- to 2-mm fragments then digested with type 2 collagenase (1.5 mg/ml), hyaluronidase (0.7 mg/ml), and type 1 DNase (0.15 mg/ml) in HBSS for 1 h at 37°C, for a total of three cycles of 1 h digestion. After each cycle, the dispersed cells were collected by filtering through a no. 80 mesh sieve and suspended in HBSS containing 1% FCS and 10 mM HEPES. Cells were suspended in HBSS, layered over a Percoll cushion, and centrifuged at 700 × *g* at room temperature for 20 min. Nucleated cells were collected from the buffer/Percoll interface. Cells enriched by Percoll density-dependent sedimentation were adjusted to a concentration of 1 × 10^6^ cells/ml in serum-free X-VIVO^TM^ 15 medium (Lonza BioWhittaker) containing 100 ng/ml recombinant human SCF (Peprotech) in 24-well plates. The culture medium was changed weekly, and wells were split when cell concentration doubled. The percentages of mast cells were assessed cytochemical by metachromatic staining with toluidine blue. Mature mast cells at ∼ 100% purity and viability are obtained by 6-8 weeks of culture, and 8- to 16-week-old mast cells were used in these experiments. HSMCs were shipped to the Huang laboratory at National Jewish Health for RNA-seq and qPCR analysis.

### HSMCs stimulations

IgE receptor cross-linking (IgECL) on HSMCs was accomplished through sensitization with 1 μg/mL chimeric human IgE anti-NP antibody (MCA333S, BIO-RAD, Hercules, CA) for 24 hours. The sensitized HSMCs were challenged with 100 ng/mL NP-HSA (N-5059, LGC, Petaluma, CA) for two additional hours prior to performing RNA-seq, ATAC-seq, or Quantitative RT-PCR analysis. For single or combined cytokine and IgECL treatments, IL-33 (50 ng/mL, 200-33, PEPROTECH, Cranbury, NJ), TSLP (50 ng/mL, 300-62, PEPROTECH, Cranbury, NJ) or IL-25 (50 ng/mL, 200-24, PEPROTECH, Cranbury, NJ) were added to HSMCs for a total of 26 hours (pre-treated for 24 hours prior to the addition of the other stimuli treatment for 2 hours).

### RNA-seq

Total RNA was extracted from HSMCs that were untreated or treated with various stimuli using the RNeasy mini kit (Qiagen, Valencia, CA) according to the manufacturer’s protocol. The RNA-seq libraries were prepared as described before [61]. Briefly, the quality of input RNA was analyzed by running the RNA sample on the 4150 TapeStation System (Agilent, CA). PolyA mRNA was isolated using the NEBNext^®^ Poly(A) mRNA magnetic isolation module (E7490, New England Biolabs, Ipswich, MA). RNA-seq libraries were obtained using the NEB-Next^®^ Ultra^TM^ II RNA library prep kit for Illumina^®^ (E7770S, New England Biolabs, Ipswich, MA). The isolated mRNA was fragmented, reverse transcribed into cDNA, end-repaired, and adapter-ligated. The ligated samples were digested by USER enzyme, amplified by PCR, and purified using AMPure-XP beads (Beckman Coulter Life Sciences, A63881, Indianapolis, IN). The quantity and quality of RNA libraries were assessed on an Agilent Technologies 2100 Bioanalyzer. Paired-ended sequencing of the RNA libraries was performed on an Illumina NovaSEQ6000 platform.

### Omni-ATAC-seq

Omni-ATAC-seq, an improved ATAC-seq protocol, was performed according to the published method [61, 62]. Briefly, 50,000 HSMCs that were untreated, treated with IL-33 alone, or IgECL with or without IL-33 were spun down and washed once with cold PBS. The cells were resuspended in 50 μl cold ATAC-RSB-lysis buffer and incubated for 3 minutes. The ATAC-RSB-lysis buffer was immediately washed out with 1 mL ATAC-RSB buffer. The cell pellet was resuspended in 50 μl transposition mix and incubated for 30 minutes at 37 °C. The reaction was stopped by adding 2.5 μl pH 8 0.5 M EDTA. The Qiagen MiniElute PCR purification kit (Qiagen) was used to purify the transposed DNA. Purified DNA was amplified using the following conditions: 72°C for 5 min, 98 °C for 30 s, and 13 cycles: 98 °C for 10s, 63 °C for 30 s, 72 °C for 1min. The amplified libraries were purified, size-selected, and the quality and quantity of libraries were assessed on 4150 TapeStation System (Agilent, CA). The pair-ended sequencing of DNA libraries was performed on an Illumina NovaSEQ6000 platform.

### RNA-seq data analysis

The RNA-seq data was analyzed as described before [61]. Briefly, the raw reads (average 20 million paired end reads, two biological replicates for each treatment) were analyzed and quality checked by FastQC. The reads were aligned to the hg38 reference genome using the Spliced Transcripts Alignment to a Reference (STAR, version 2.4.0.1) software. Reads (FPKM) were assembled into reference transcripts and counted using Cufflinks (version 2.2.1). The average reads from two biological samples were calculated using Cuffmerge (version 1.0.0). The differential gene expression between the resting and stimulated samples was analyzed using Cuffdiff (version 2.2.1). K-means clustering was performed using pheatmap (version 1.0.12). The optimal numbers of K in the RNA-seq datasets were determined using R packages Tidyverse (version 1.3.1), Cluster (version 2.1.0), Factoextra (version 1.0.7), and GridExtra (2.3). Genes in clusters were extracted using pheatmap (version 1.0.12).

### Omni-ATAC-seq data analysis

The Omni-ATAC-seq data was analyzed as described before [61]. Briefly, raw sequencing reads (average 80 million paired-end reads, two biological replicates for each treatment) were aligned to the hg38 reference genome using Bowtie2 with very-sensitive and -x 2000 parameters. The read alignments were filtered using SAMtools to remove mitochondrial genome and PCR duplicates. Narrow peaks were called by MACS2 with the q-value cut-off of 0.05, and the sequencing data was displayed using Integrative Genomics Viewer (IGV).

### Quantitative RT-PCR

For mRNA expression analysis, the HSMCs were untreated, treated with IL-33 or IgECL alone or in combination. Total RNA from HSMCs was isolated with TRIzol^TM^ Reagent according to the manufacturer’s instructions. cDNA was synthesized by reverse transcription. Quantitative PCR was performed in a QuantStudio 7 Flex Real-Time PCR System (ThermoFisher, MA). The sequences of qPCR primers are listed in Table S6. Relative mRNA amounts were calculated as follows: Relative mRNA amount = 2^[Ct(*Sample*)-Ct(*HPRT*)]^.

### ELISA measurement of chemokine and cytokine productions

HSMCs were pretreated with IL-33 (50 ng/mL, 200-33, PEPROTECH, Cranbury, NJ), or sensitized with 1 mg/mL chimeric human IgE anti-NP antibody (MCA333S, BIO-RAD, Hercules, CA), or in combination for 24 hours. Then washed before stimulation with 100 ng/mL NP-HSA (N-5059, LGC, Petaluma, CA) or 50 ng/mL IL-33, or in combination for additional 24 hours. Untreated HSMCs were included as a control. The amounts of CXCL8 (88-8086-22, Thermo Fisher Scientific), IL1B (88-7261-22, Thermo Fisher Scientific), and CXCL2 (900-M120, Peprotech) protein in the supernatants were measured by ELISA.

### DAVID GO analysis

Gene ontology enrichment analysis was performed on genes that responded to the synergistic stimulations of IgECL plus IL-33 by using the Database for Annotation, Visualization, and Integrated Discovery (DAVID, version 6.8). The top-ranked genes in each enriched category were selected for heatmap representations using an Excel spreadsheet.

### Assigning regulatory regions to genes

Binding and Expression Target Analysis (BETA) minus (Version 1.0.7) used to calculate the regulatory potential score and assign the regulatory regions identified by Omi-ATAC-seq to their putative target genes [51] with the following setting: Reference genome, Human: GRCh38; Distance: within 100 kb upstream and 100 kb downstream from the transcriptional start site of the gene.

### DiffBind

For analysis of activation-induced accessible regions, Omni-ATAC-seq reads of HSMCs samples that were treated with IL-33 and IgECL were compared with those of HSMCs samples treated with IL-33 or IgECL using the R package DiffBind (version 2.10.0) [63]. The number of upregulated peaks in the synergy genes were calculated by adding the number of short peaks within 100 kb upstream and 100 kb downstream from TSS were significantly increased in response to the combined IL-33 priming and antigenic stimulation relative to IL-33 or IgECL.

### TF binding motif enrichment analysis

Multiple Em for Motif Elicitation (MEME)-ChIP performs comprehensive motif analysis on large sets of sequences such as those identified by ChIP-seq or CLIP-seq experiments [52]. MEME-ChIP was used to identify the TF binding motifs enriched in accessible regions. To identify the TF binding motifs enriched in the induced accessible regions, BED files of IgE-stimulated and IgE+IL-33-stimulated were uploaded to the Find Individual Motif Occurrences (FIMO) tool of MEME (FIMO) [64]. The number of transcription factor binding sites in the increased narrow peaks of the synergy genes was calculated by adding the number of transcription factor binding sites within the narrow peaks. The densities of transcription factor binding sites in the increased narrow peaks were normalized by the sizes of narrow peaks and expressed as the number of binding sites per kilobase.

### Data availability

All raw and processed sequencing data generated in this study have been submitted to the NCBI Gene Expression Omnibus (GEO; https://www.ncbi.nlm.nih.gov/geo/) under accession number GSE196895. The RNA-seq data can be found in the GEO database: “GSE196862”; The ATAC-seq data can be found in the GEO database: “GSE196883”.

### Statistical analysis

One-tail Fisher Exact probability value was used for GO analysis. Statistical differences among the single and combined treatments were analyzed using two-way ANOVA [29]. Identifying statistically significantly differentially bound sites in DiffBind used specifical statistical methods built in the Bioconductor packages EdgeR [65] and DEseq2 [66].

## Abbreviations

HSMCs: Human skin mast cells
RNA-seq: RNA sequencing
ATAC-seq: Transposase-accessible chromatin sequencing
IgECL: IgE cross-linking
TSLP: Thymic stromal lymphopoietin
qRT-PCR: quantitative reverse transcription-PCR
NF-κB: Nuclear factor kappa-light-chain-enhancer of activated B cells
AP-1: Activator protein 1
GABPA: GA-binding protein alpha chain
RAP-1: Ras-related protein 1
NFAT: nuclear factor of activated T-cells
MAP: mitogen-activated protein
NP: Nitrophenyl-hapten
GO: gene ontology
FPKM: Fragments Per Kilobase of transcript per Million mapped reads
IGV: Integrative Genomics Viewer
DAVID: Database for Annotation, Visualization, and Integrated Discovery
BETA: Binding and Expression Target Analysis
MEME: Multiple Em for Motif Elicitation
TSS: Transcription start site

## Declarations

### Ethics approval and consent to participate

The study was performed in accordance with the Declaration of Helsinki on research involving humans. The skin tissues used to obtain the mast cells for this study were purchased from the Cooperative Human Tissue Network of the National Cancer Institute, and informed consent was obtained from all participants. This study was reviewed by the Institutional Review Board (IRB) of the University of South Carolina. The IRB of the University of South Carolina has determined that the referenced study (protocol number Pro00020620) meets the Not Human Subject criteria set forth as defined by the Code of Federal Regulations (45 CFR 46) as the specimens and/or private information/data were not collected specifically for the current research project through an interaction/intervention with living individuals and the investigators including collaborators on the proposed research cannot readily ascertain the identity of the individuals to whom the coded private information or specimens pertain. All methods and experiments were performed in accordance with relevant guidelines and regulations.

### Consent for publication

Not applicable

### Availability of data and materials

The RNA-seq datasets and ATAC-seq datasets generated in this study have been submitted to the NCBI Gene Expression Omnibus (GEO; https://www.ncbi.nlm.nih.gov/geo/) under accession number GSE196895.

### Authors’ contributions

Conceptualization, J.G. and H.H.; methodology, J.G., Y.L., G.G., C.A.O. and H.H.; investigation, J.G., Y.L., X.G., Z.M., D.Z. and Y.H.; visualization, J.G. and Y.L.; funding acquisition, H.H., C.A.O. and G.G.; supervision, H.H., C.A.O. and G.G.; writing-original draft, J.G. and H.H.; writing-review & editing, J.G., C.A.O. and H.H.

## Acknowledgements

We thank lab members for the thoughtful discussions. We are grateful to Dr. Bifeng Gao and the staff of the Genomics Shared Resource Facility at the University of Colorado Cancer Center for Next-Generation Sequencing.

