## Supplementary material for "IL-33 priming and antigenic stimulation synergistically promote the transcription of proinflammatory cytokine and chemokine genes in human skin mast cells": Figure S1 and Figure S2

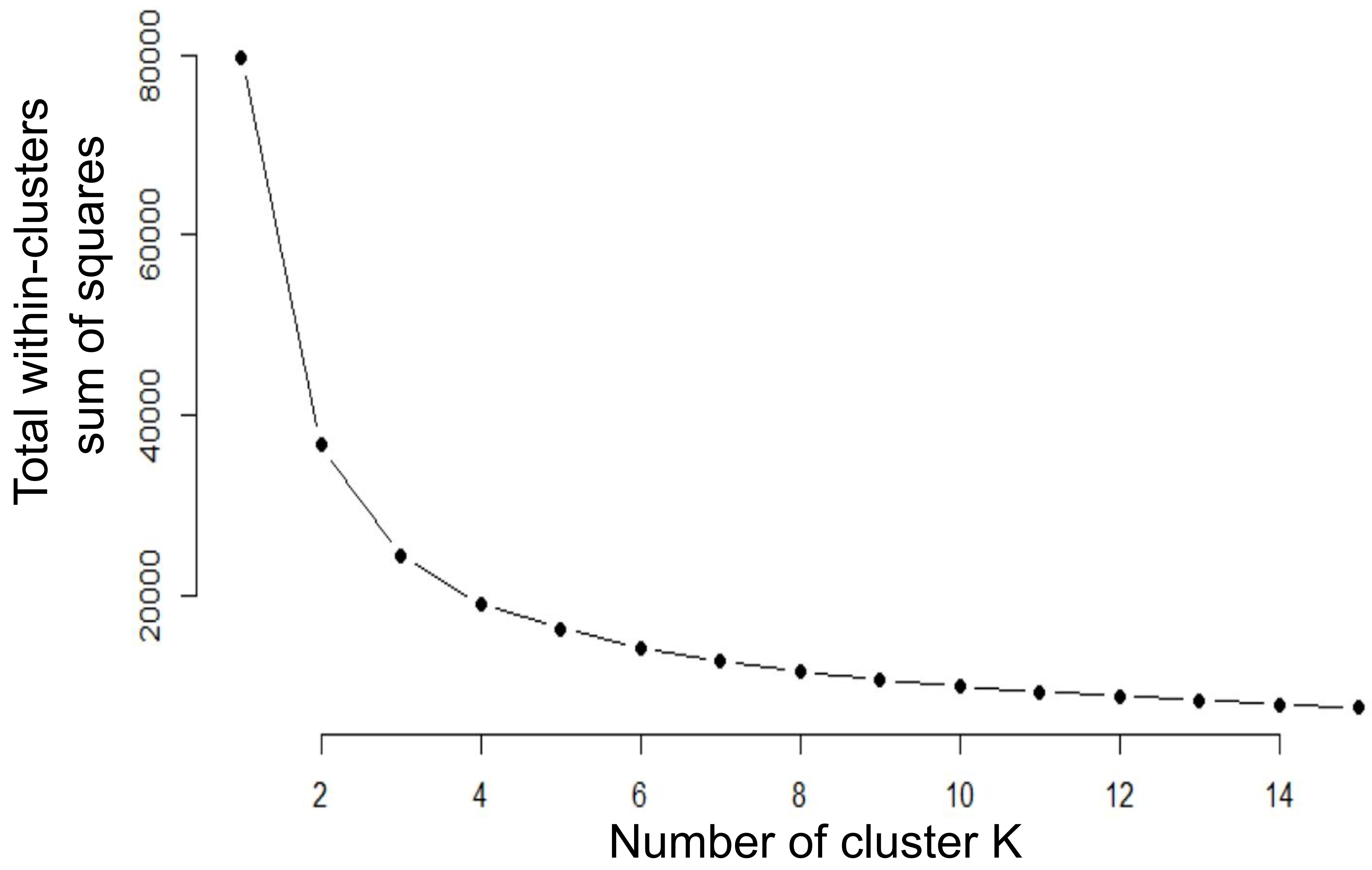


**Figure S1.** K-means clustering elbow plots. The optimal numbers of K in the RNA-seq datasets (two biological samples) were determined using R packages Tidyverse (version 1.3.1). The within cluster sum of squares decrease with the increment of clusters number, and the optimal number of clustering was selected at the last one significantly reduced the within cluster sum of squares (at the inflection point of the curve). The optimal clustering was at number of six for the RNA-seq datasets.

### Figure S2

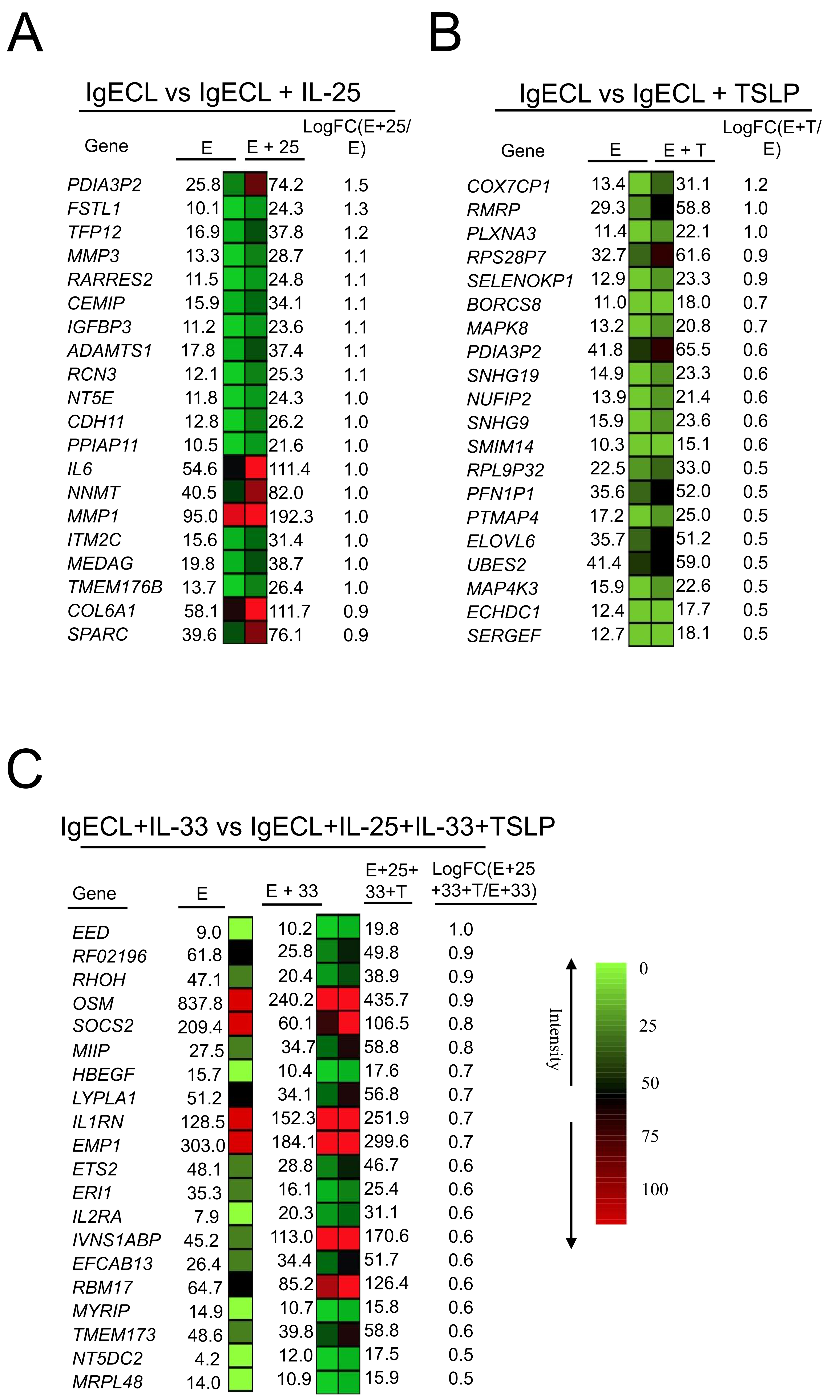


**Figure S2**. Differential mRNA expression analysis of RNA-seq prepared from HSMCs received various combined treatments. Heatmap representations of top ranked genes after the combined IgECL+IL-25 treatment (A), IgECL+TSLP treatment (B) or IgECL+IL-33+IL-25+TSLP treatment (C). Legends: E, IgECL; E+25, IgECL+IL-25 treatment; E+T, IgECL+TSLP treatment; E+33, IgECL+IL-33 treatment; E+25+33+T, IgECL+IL-25+IL-33+TSLP treatment; LogFC, log2 fold change. The numbers indicate RNA reads (RPKM). Data A-C represent two biological samples.
